# Red blood cell-derived extracellular vesicles with miR-204 mimic loading for pediatric neuroblastoma treatment

**DOI:** 10.1101/2023.05.29.542704

**Authors:** Wararat Chiangjong, Jirawan Panachan, Sujitra Keadsanti, David S. Newburg, Ardythe L. Morrow, Suradej Hongeng, Somchai Chutipongtanate

**Author notes:** Correspondence (W.C.);, (S.C.).

## Abstract

Neuroblastoma (NB) is the most common extracranial solid tumor in pediatric population with a high degree of heterogeneity in clinical outcomes, ranging from spontaneous remission to rapid progression and death. Upregulation of a tumor suppressor miR-204 in patient-derived neuroblastoma tumors was associated with good prognosis independent of known risk factors. While miR-204 is recognized as a therapeutic candidate, its delivery was unavailable. This study aimed to develop red blood cell-derived extracellular vesicles (RBC-EVs) as the miR-204 carrier and evaluate the inhibitory activity against neuroblastoma cell lines and spheroids. MiR-204 mimics were loaded into RBC-EVs (RBC-EV^miR-204^) by electroporation with the optimized parameters of 250 V, 20 ms, 10 pulsing times. RBC-EV^miR-204^, but not the native RBC-EVs, could inhibit cell viability, migration and spheroid formation and growth of MYCN-amp and MYCN non-amplification (MYCN-NA) NB cells, even though the suppressive effects were more preferable in MYCN-amp NB. For the mechanistic insight, SWATH-proteomics suggested that RBC-EV^miR-204^ induced dysregulation of ribosomal proteins and alterations in RNA metabolism, leading to inhibiting neuroblastoma progression. This study developed RBC-EV^miR-204^ as an alternative/adjunct therapy of pediatric neuroblastoma. The therapeutic efficacy of RBC-EV^miR-204^ should be further investigated in preclinical models and clinical studies.

## Introduction

Neuroblastoma (NB) arising from the peripheral sympathetic nervous system is the most common extracranial solid tumor in the pediatric population, [1,2]. NB accounts for 15% of all childhood cancer-associated mortalities [1,2]. A high degree of NB heterogeneity leads to a wide array of clinical outcomes, ranging from spontaneous remission to rapid progression and death [3,4]. Thus, NB treatment currently follows a consensus approach for pretreatment risk classification as proposed by the International Neuroblastoma Risk Group [5] classification system, including clinical features, tumor histology, and molecular markers including *MYCN* amplification (*MYCN*-amp) and 11q aberration [1,2,5,6]. Radiotherapy, chemotherapy, and observation treatment can completely eliminate the MS stage of neuroblastoma with >90% cure rate [2,7]. Although the high-risk stage of NB patients with poor biological characteristics was treated with high-intensity treatment strategies including 5-6 rounds of surgery, high-dose combined radiation, chemotherapy, autologous stem cell transplantation, and anti-GD2 immunotherapy, the outcome remained poor with a cure rate of < 50% and clinically significant side effects [2,8-11].

MicroRNA-204 (miR-204) is recognized as a therapeutic candidate for NB. MiR-204 is known as a tumor suppressor, targeting multiple oncogenes involved in tumorigenesis and progression, i.e., *BDNF, MEIS1, FOXC1, NUAK1* and *RAB22A* [12-17]. MiR-204 may exhibit dual functions of tumor-suppressor and oncogene in prostrate and breast cancers [18]. In pediatric neuroblastoma, Ryan et al. [19] revealed a significant low expression of miR-204 in patients with high-risk factors, including *MYCN-*amp. Multivariate Cox proportional hazard regression analysis showed upregulation of miR-204 (greater than median expression) in neuroblastoma tumor tissues was significantly associated with improved event free survival (HR 0.38, 95% CI 0.17-0.91; p=0.021) independent of age, *MYCN*-amp, 11q deletion, and INSS stages [19]. MiR-204 also targets several oncogenes in NB. miR-204 binds *MYCN* mRNA to repress N-Myc protein expression, and conversely, *MYCN* mRNA binds to the miR-204 promoter to repress miR-204 transcription [20]. *BCL2* (anti-apoptotic gene) and *NTRK2* (oncogene) activation is associated with poor NB patient survival; MiR-204 targets the 3’ UTR of each of these genes, resulting in enhanced chemosensitivity of neuroblastoma cells to cisplatin and etoposide [19]. The *PHOX2B* 3′UTR gene is overexpressed in NB tumor samples and NB cell lines. MiR-204 binding in the two sites proximal and distal within the most conserved region of the PHOX2B 3′UTR gene, is associated with tumor suppression [21].

The introduction of miRNA mimics systemically may induce serious immune-mediated adverse events e.g., the early termination of phase I trial of miR-34 encapsulated liposomal nanoparticles for advanced solid tumors [22]. Therefore, developing an effective and compatible delivery system is the greatest challenge of miRNA therapy [22,23]. The blood-brain and tumor barrier are also a major obstacle. Safe and precise nanotechnology-based delivery systems, i.e., nanosized gold, carbon and silica particles with complexation, encapsulation, and conjugation-based approaches, were developed to deliver miRNAs into NB cells [24,25]. Specific targeting nanoparticles were created to penetrate the blood-brain and tumor barrier such as 2-[3-[5-amino-1-carboxypentyl]-ureido]-pentanedioic acid (Acupa) and cyclic TT1 (cT) co-functionalized nanoparticles (A-NPs-cT) whose specificity is to prostate-specific membrane antigen in breast cancer brain metastases [26]. Both bioengineering nanoparticles and specificity of molecular targets for crossing the blood-brain barrier are impediments to this approach.

To overcome these engineering and specific target searching impediments, biological materials such as red blood cell extracellular vesicles (RBC-EVs) can be used as an efficient RNA drug delivery system [27,28]. Drug-loaded RBC-EVs have low side effects on normal cells, are stable and remain intact even after multiple freeze-thaw cycles, and during storage at - 80°C without affecting the moiety, uptake, and genetic material loading capacity [28,29]. They can cross the blood-brain barrier without any specific target on their surface, escape the immune system via CD47 to prevent erythrocyte uptake by macrophages, and remain in blood circulation for 44 min [30,31]. Thus, RBC-EV properties can overcome many of obstacles intrinsic to drug delivery systems, expediting internalization into cancer cells, thereby delivering therapeutic drugs to eradicate cancer cells.

This study was designed to develop and test miR-204-loaded RBC-EVs (RBC-EV^miR-204^) as an alternative strategy for NB treatment. MiR-204 mimic was loaded into RBC-EVs using electroporation, a well-known method for passive packaging of all kinds of cargos into EVs [32], following by examining anti-neuroblastoma effects against several NB cell lines and investigating RBC-EV^miR-204^ mechanisms of inhibition in NB cells using SWATH-proteomics.

## 2. Materials and Methods

### 2.1. Cell cultivation

Neuroblastoma cell lines SK-N-BE2 (ATCC: CRL-2271), SH-SY5Y (ATCC: CRL-2266), SK-N-AS (ATCC: CRL-2137), SK-N-DZ (ATCC: CRL-2149) and SK-N-SH (ATCC: HTB-11) and cell lines A549 (ATCC: CRM-CCL-185), MDA-MB-231 (ATCC: CRM-HTB-26), HepG2 (ATCC: HB-8065), HT29 (ATCC: HTB-38) and Jurkat (ATCC: CRL-2899) were purchased from American Type Culture Collection. SH-SY5Y, SK-N-AS, SK-N-DZ, A549, HepG2 and MDA-MB-231 cells were maintained in DMEM-HG supplemented with 10% FBS (Gibco, Thermo Fisher Scientific, MA, USA) and 1× penicillin/streptomycin (Gibco). SK-N-BE2 and HT-29 cells were grown in DMEM/F12 supplemented with 10% FBS and 1× penicillin/streptomycin. Jurkat cells were cultured in RPMI-1640 supplemented with 10% FBS and 1× penicillin/streptomycin. SK-N-SH cells were maintained in DMEM supplemented with 10% FBS and 1× penicillin/streptomycin. All cells were cultured under a humidified atmosphere with 5% CO_2_ at 37°C for 24-48 h.

Activated T cells were derived from 3 healthy donors under a protocol approved by Human Research Ethics Committee, Faculty of Medicine Ramathibodi Hospital, Mahidol University, based on the Declaration of Helsinki (COA. MURA2020/2010). After informed consent, 10 ml EDTA whole blood was collected from participants, diluted in PBS at a 1:1 ratio, and then carefully layered onto the Ficoll-Paque solution (Robbins Scientific Cooperation, Norway) at a 2:1 ratio (diluted blood: Ficoll-Paque solution). The tube containing layer solution was centrifuged at 400 g, 20°C for 15 min with no break. The human peripheral blood mononuclear cell (PBMC) layer was transferred into a new centrifuge tube, and RBCs were collected for RBC-EV production. PBMCs were washed with PBS twice, and cultured in an anti-CD3 (MACS, Miltenyi Biotec) and anti-CD28 (MACS, Miltenyi Biotec) coated plate in 1 µg of each antibody in 1 ml PBS per 24-well plate for 2 h at room temperature. For culturing, PBMCs were added into the coated well which had been washed with PBS once, and cultured in RPMI-1640 supplemented with 100U/ml IL-2 (PeproTech, Rocky Hill, NJ, USA), 10% FBS and 1x penicillin/streptomycin for 3 days at 37°C under a humidified atmosphere with 5% CO_2_ incubator to obtain T cell blasts. The culture medium was replaced every other day with fresh media containing 100U/ml IL-2.

### 2.2. RBC-EV production, isolation and characterization

Calcium ionophore was used to stimulate EV production of RBCs [27]. After removing plasma and WBCs, RBCs were washed thrice in HEPES. Packed RBCs (500 µl) were resuspended in 1 mL of HEPES buffered physiological solution (HPS) containing 145 mM NaCl, 7.5 mM KCl, 10 mM glucose, 10 mM HEPES, pH 7.4. The 1.5 ml resuspended RBCs were cultured in 50 ml HPS containing 2 mM CaCl_2_ and 2 µM calcium ionophore-4-bromo-A23187 (C7522, Sigma-Aldrich, Merck KGaA, Darmstadt, Germany) in T75 flasks in a 5% CO_2_ incubator at 37°C for 16 h. The supernatants were collected in 50 mL tubes and centrifuged at 3000 g for 15 min twice to remove the RBC pellet. The supernatant was transferred to 100 kDa cut-off ultrafiltration column (Vivaspin20, Cytiva, Cytiva Sweden AB, Uppsala, Sweden) and centrifuged at 4000 g for 15-45 min until the UF-concentrated solution of 0.5 mL remained. EVs were then isolated from the soluble protein contaminants by qEV size exclusion chromatography (qEVoriginal/35 nm Legacy, IZON Science Ltd., Christchurch, New Zealand) as previously described [33,34]. Briefly, the qEV column was equilibrated with 3 column volumes (30 mL) of HPS and then loaded with the UF-concentrated solution (0.5 mL). After the fraction solution was completely absorbed in the beads within the qEV column, the column was washed with 3 ml HPS and the wash discarded. Thereafter, 2.5 mL HPS was added into the column to collect the EV fraction. An additional 5.5 mL HPS washed the proteins and other contaminants from the column, and was discarded. The RBC-EV fractions were concentrated using 3 kDa cut-off column and stored at -80°C until use. The RBC-EVs were confirmed by morphology and size distribution by TEM and nanosight tracking analysis (NTA), and the EV marker TSG101 protein measured by Western blot analysis. Mass spectrometric proteomics confirmed evidence of RBC-EV protein. Pooled RBCs and pooled RBCEVs were lysed in reducing buffer containing 62.5 mM Tris-HCl, pH 6.8, 10%(v/v) glycerol, 2%(w/v) SDS and 2.5%(v/v) β-mercaptoethanol. After complete lysis, the protein content was measured by Bradford’s assay. Twenty micrograms of total proteins were then reduced, alkylated, and digested with trypsin. One microgram samples were analyzed by LC-MS/MS (n=10 technical replications for IDA). Ten IDA .*wiff* files (10 files of each RBC and RBCEV preparation) were searched against human with <1%FDR.

### 2.3. Effect of RBC-EVs on various cancer cell types

Approximately 2×10^4^ cells of eight cancer cell lines (SK-N-BE2, SH-SY5Y, SK-N-AS, SK-N-SH, A549, MDA-MB-231, HepG2, HT-29 and Jurkat cells), one normal human fibroblast line (HEK-293T) and one primary cells (peripheral blood T cells) were seeded into each well of a 96-well plate and then incubated at 37°C overnight. The cultured media in each well was replaced with fresh complete medium containing 0, 100, 1000 and 10000 RBC-EV particles/cell (n=3) in 100 µL/well. After a 3-day incubation, cell viability was measured by the MTT assay (Cell proliferation kit, Roche Diagnostics Deutschland GmbH, Munich, Germany). Briefly, 10 µL MTT labeling reagent was added into each well and then incubated at 37°C for 4 h.

Thereafter, 100 µL solubilization solution was added into each well and then incubated at 37°C overnight. Colorimetric detection was at 570 nm. The optical density of the blank wells containing the complete medium without cells was subtracted from the wells containing cells. The % cell viability was calculated as: % cell viability= OD treatment/OD non-treatment ×100). This experiment was performed in three independent experiments per cell line.

### 2.4. MiRNA loading into RBC-EVs

One hundred picomoles of miR-204 (5’-UUCCCUUUGUCAUCCUAUGCCU-3’; with or without 5’FAM tag) or miRNA mimic negative control (miRNC; 5’-CGGUACGAUCGCGGCGGGAUAUC-3’) (Macrogen, Inc., Seoul, South Korea) were mixed with 1×10^9^ RBC-EV particles and adjusted to 100 µL final volume. The mixture was transferred to Gene Pulser/MicroPulser Electroporation Cuvettes (Biorad). The electroporation conditions varied voltages with a constant pulsing time (50V 20ms, 100V 20ms, 250V 20ms, 400V 20ms), 750V 1ms, and 250V with varied pulsing time 5, 10, and 20ms. The electroporated mixture was transferred into a 1.5 mL tube to which is added PKH26 dye (0.4 µL) (PKH26 Red Fluorescent Cell Linker Kit for General Cell Membrane Labeling, Sigma) mixed with 100 µL diluent C. After 5 min incubation at room temperature, the staining reaction was stopped by adding 200 µL FBS (an equal volume of the staining solution). The reaction was transferred into SK-N-BE2 cells and then incubated at 37°C for 2 days. Thereafter, the treated cells were stained with Hoechst33342 dye for 30 min at 37°C before washing with PBS, fixing with 4% paraformaldehyde/PBS, mounting in 20% glycerol/PBS and the images captured by a confocal microscope (Nikon).

### 2.5. Effects of RBC-EV^miR204^ on NB cells

#### 2.5.1. Cell viability

Approximately 2×10^4^ SK-N-BE2 or SH-SY5Y NB cells in DMEM:F12 or DMEM-high glucose medium containing 10% FBS (Gibco), respectively were seeded into a well of 96-well plate overnight. One milliliter of basal medium containing RBC-EV^miR204^ or RBC-EV^miRNC^ (1000 particles/cell) (n=3) were then added. The native RBC-EVs (1000 particles/cell) and non-treated cells served as the controls. After a 2-day incubation, cell viability was measured by MTT assay (Cell proliferation kit, Roch2.2e). Three independent experiments were performed per cell line.

#### 2.5.2. Cell migration

A wound scratch assay was a surrogate measure to evaluate NB cell migration potential as representative of metastasis. Approximately 1.5×10^6^ SK-N-BE2 or SH-SY5Y cells in DMEM:F12 or DMEM-high glucose medium containing 10% FBS and 1× penicillin/streptomycin were cultured in wells of 12-well plate overnight. A wound was created by scratching using a 200-µL pipette tip and washing unattached cells with PBS once. Then, 1 mL basal medium containing RBC-EV^miR-204^ or RBC-EV^miRNC^ (1000 particles/cell) were added into each well (n=3) and then the wound images were captured at 0 and 48 h. The native RBC-EVs and non-treatment served as the controls. The wound gap was measured by using ImageJ software (www.imagej.nih.gov). Three independent experiments were performed per cell line.

#### 2.5.3. Spheroid forming and growing assay

Approximately 3×10^3^ SK-N-BE2, SH-SY5Y, SK-N-DZ, and SK-H-SH cells in DMEM:F12, DMEM-high glucose medium and DMEM containing 1% knock-out serum replacement (Gibco), respectively, were seeded into wells of 96-well clear, ultra-low binding, U-shaped bottom, Corning spheroid microplates (Corning®, Arizona, USA) in the presence of RBC-EV^miR-204^ or RBC-EV^miRNC^ (1000 particles/cell) (n=3). The native RBC-EVs and non-treatment served as controls. Both cell forming and growing were measured at 0, 24 and 48 h after treatment. The spheroid diameter was measured by ImageJ software. This experiment was performed in three independent experiments per cell line.

### 2.6. Quantitative mRNA expression of MYCN genes

Approximately 1.5×10^6^ SK-N-BE2 in 1 mL DMEM:F12 or SH-SY5Y cells in 1 mL DMEM-high glucose medium containing 10% FBS and 1× penicillin/streptomycin were cultured in wells of 12-well plates overnight. One milliliter of the basal medium containing RBC-EV^miR204^ or RBC-EV^miRNC^ (1000 particles/cell) were added. The native RBC-EVs and non-treatment served as the controls. After 48 h in culture (n=3), cells were collected in PBS and washed twice with PBS by centrifugation at 400 g for 5 min at RT. After removing PBS from the cell pellets, 1 mL TRIzol^®^ reagent (Invitrogen, Carlsbad, CA, USA) was added into the cell pellet to extract total RNA. Quantity and quality of RNA were determined at OD 260 nm and 260/280 ratio, respectively, by MULTISKAN GO (Thermo SCIENTIFIC). One microgram of RNA was converted to cDNA using ImProm-II™ Reverse Transcription System (PROMEGA, Madison, WI, USA) following the manufacturer’s instructions performed on T100™ Thermal Cycler (BIO-RAD). Reverse transcription protocol was: annealing at 25°C for 10 min, extended the first strand at 42°C for 1 h, and heat-inactivated the reverse transcriptase at 70°C for 15 min. Quantitative PCR was performed using *MYCN* and β-actin primers (Biotechrabbit, biotechrabbit GmbH, Berlin, Germany) as following; *MYCN* forward and reverse primers: 5’CACAAGGCCCTCAGTACCTC 3’ and 5’ ATGACACTCTTGAGCGGACG 3’; β-actin forward and reverse primers: 5’ AGAGCTACGAGCTGCCTGAC 3’ and 5’ AGCACTGTGTTGGCGTACAG 3’. PCR amplification condition was 95°C for 3 min, and then 50 cycles of 95°C for 15s and 65°C for 30s. *MYCN* mRNA intensity was normalized with β-actin mRNA intensity as well as relative to the non-treatment group. Three independent experiments were performed per cell line.

### 2.7. SWATH-proteomics

SWATH-proteomic and protein bioinformatic analyses were performed as described previously [35-37]. Approximately 1.5×10^6^ SK-N-BE2 cell lines in 1 mL DMEM:F12 or SH-SY5Y in 1 mL DMEM-high glucose medium were cultured in wells of 12-well plates in the presence of RBC-EV^miR204^ or RBC-EV^miRNC^ at 1000 particles/cell for 48 h (n=3). The native RBC-EVs and non-treated cells served as the controls. Cells were collected and washed with PBS twice by centrifugation at 200 g for 5 min at RT. The reducing buffer containing 62.5 mM Tris-HCl, pH 6.8, 10%(v/v) glycerol, 2%(w/v) SDS and 2.5%(v/v) β-mercaptoethanol was added into the cell pellet and then mixed by Vortex until the pellet dissolved. The protein was measured by using the Bradford assay. Forty micrograms of protein were then reduced by DTT and alkylated by IAA before tryptic digestion. The peptides were desalted on a C18 stage tip before dissolving in 0.1% formic acid and injecting 1 µg into Eksigent nanoLC ultra nanoflow high-performance liquid chromatography coupled with Triple-TOF 6600+ (ABSciex, Toronto, Canada) mass spectrometry (installed at Faculty of Medicine Ramathibodi Hospital, Bangkok, Thailand) in information-dependent acquisition (IDA) and data independent acquisition modes. The peptides were loaded onto a C18 trap column (Nano Trap RP-1, 3 μm 120 Å, 10 mm × 0.075 mm; Phenomenex, CA, USA) at a flow rate of 3 μl/min in 0.1% formic acid for 10 min to desalt and concentrated the sample before separating with a C18 analytical column (bioZen Peptide Polar C18 nanocolumn, 75 μm × 15 cm, C18 particle sizes of 3 μm, 120 Å; Phenomenex) with a gradient 3-30% acetonitrile (ACN)/0.1%formic acid for 60 min, following 30-40%ACN/0.1% formic acid for 10 min, 40-80% ACN/0.1%formic acid for 2 min, isocratic 80%ACN/0.1%formic acid for 6 min, 80-3%ACN/0.1%formic acid for 2 min, and isocratic 3%ACN/0.1%formic acid for 25 min. The eluate was ionized and sprayed into the mass spectrometer using OptiFlow Turbo V Source (ABSciex). Ion source gas 1 (GS1), ion source gas 2 (GS2), and curtain gas were set at 19, 0, and 25 vendor arbitrary units, respectively. The interface heater temperature and ion spray voltage were kept at 150°C and at 3.3 kV, respectively.

Mass spectrometry was operated in positive ion mode set for 3,500 cycles for 105 min gradient elution. Each cycle performed 1 time of flight (TOF) scan (250 ms accumulation time, 350–1250 m/z window with a charge state of +2) followed by IDA of the 100 most intense ions, while the minimum MS signal was set to 150 counts. MS/MS scan was operated in a high sensitivity mode with 50 ms accumulation time and 50 ppm mass tolerance. Former MS/MS candidate ions were excluded for a period of 12 sec after its first occurrence to reduce the redundancy of identified peptides. DIA mode was performed in a range of 350 to 1500 m/z using a predefined mass window of 7-m/z with the overlap of 1-m/z for 157 transmissible windows.

MS scan was set at 2,044 cycles, where each cycle performs 1 TOF-MS scan type (50 ms accumulation time across 100–1500 precursor mass range) acquired in every cycle for a total cycle time of 3.08 sec. MS spectra of 100–1500 m/z were collected with an accumulation time of 96 ms per SWATH window width. Resolution for MS1 scan was 35,000 and SWATH-MS2 was 30,000. Rolling collision energy mode with collision energy spread of 15 eV was applied. The IDA and DIA data (.*wiff*) were recorded by Analyst-TF v.1.8 software (ABSciex).

A total of 12 wiff files of IDA experiments per cell line were combined and searched using Protein Pilot v.5.0.2.0 software (ABSciex) against the Swiss-Prot database (UniProtKB 2022_01) Homo sapiens (20,385 proteins in database) with the searching parameters as follows; alkylation on cysteine by iodoacetamide, trypsin enzymatic digestion, 1 missed cleavage allowed, monoisotopic mass, and 1% false discovery rate. The group file (Protein Pilot search result) was loaded into SWATH Acquisition MicroApp v.2.0.1.2133 in PeakView software v.2.2 (Sciex) to generate a spectral library. The maximum number of proteins was set as the number of proteins identified at 1% global FDR from fit. RT alignment was performed by the high abundance endogenous peptides covering the chromatographic range. SWATH data extraction of 12 DIA files per cell line was performed by SWATH Acquisition MicroApp (ABSciex) using the following parameters; 10-min extraction window, 25 peptides/protein, 6 transitions/peptide, excluding shared peptides, >99% peptide confidence, <1% FDR, and 20 ppm XIC width.

SWATH extraction data, including the identities and quantities of peptides and proteins, was exported into Marker view (ABSciex). The protein area was normalized by multiple linear regression normalization and calculated the differential protein expression using ANOVA and fold-change of protein area. The protein interaction pathway was predicted by STRING (https://string-db.org).

#### 3. Results

### 3.1. Preliminary data on the effect of free miR-204 mimics against NB cell viability

A preliminary study on the effect of free miRNA was performed in *MYCN*-amp SK-N-BE2 and *MYCN*-non-amplification (*MYCN*-NA) SH-SY5Y NB cells (**Figures S1-S5**). MiRNAs were introduced into cells by the optimized electroporation conditions (**Figures S1 and S2**) and the cell viability was measured at 24, 48 and 72 h (**Figure S3**). The anti-neuroblastoma effect of the free miR-204 introduced by electroporation was more pronounced in *MYCN*-amp SK-N-BE2 NB cells compared to *MYCN*-NA SH-SY5Y cells. A common lipid nanoparticle-drug delivery system, lipofectamine, was used to transfer miRNAs into the cells. The results showed that the lipofectamine-miR-204 complex exhibited higher cytotoxicity in SK-N-BE2 cells than in SH-SY5Y cells (**Figure S4**). However, lipofectamine without miRNA also exhibited significant cytotoxicity (**Figure S4**), consistent with previously reported systemic side effects of miR-34a-loaded liposomal nanoparticles in a phase I clinical trial for treating advanced solid tumors [22]. Therefore, lipid nanoparticles as the delivery system for miRNA-based therapy may not be fully compatible with clinical applications. Finally, without miRNA introduction by electroporation/lipofectamine, the free miR-204 apparently had little or no cytotoxic effect against *MYCN*-amp and *MYCN*-NA NB cells (**Figure S5**). Taken together, these preliminary findings supported further testing of RBC-EVs as effective and biocompatible miR-204 nanocarriers for NB treatment.

### 3.2. The basal effect of isolated RBC-EV on multiple cancer and normal cells

RBCs were stimulated by calcium ionophore to release RBC-EVs into culture medium [27]. RBC-EVs were then isolated by the combination of centrifugation, ultrafiltration and qEV size exclusion chromatography, followed by validation of EV presence by NTA particle size measurement, TEM particle morphological analysis and specific EV marker detection using Western blotting with anti-TSG101 antibody. As a result, RBC-EVs produced from 3 individual RBCs (three technical replicates per biological sample) were consistent in the particle size with the average diameter of 132±2 nm, 126±11 nm, and 129±19 nm (**Figure 1a**). TEM with negative staining revealed RBC-EV morphology of round-shaped nanoscale vesicles (**Figure 1b**).

**Figure 1.**
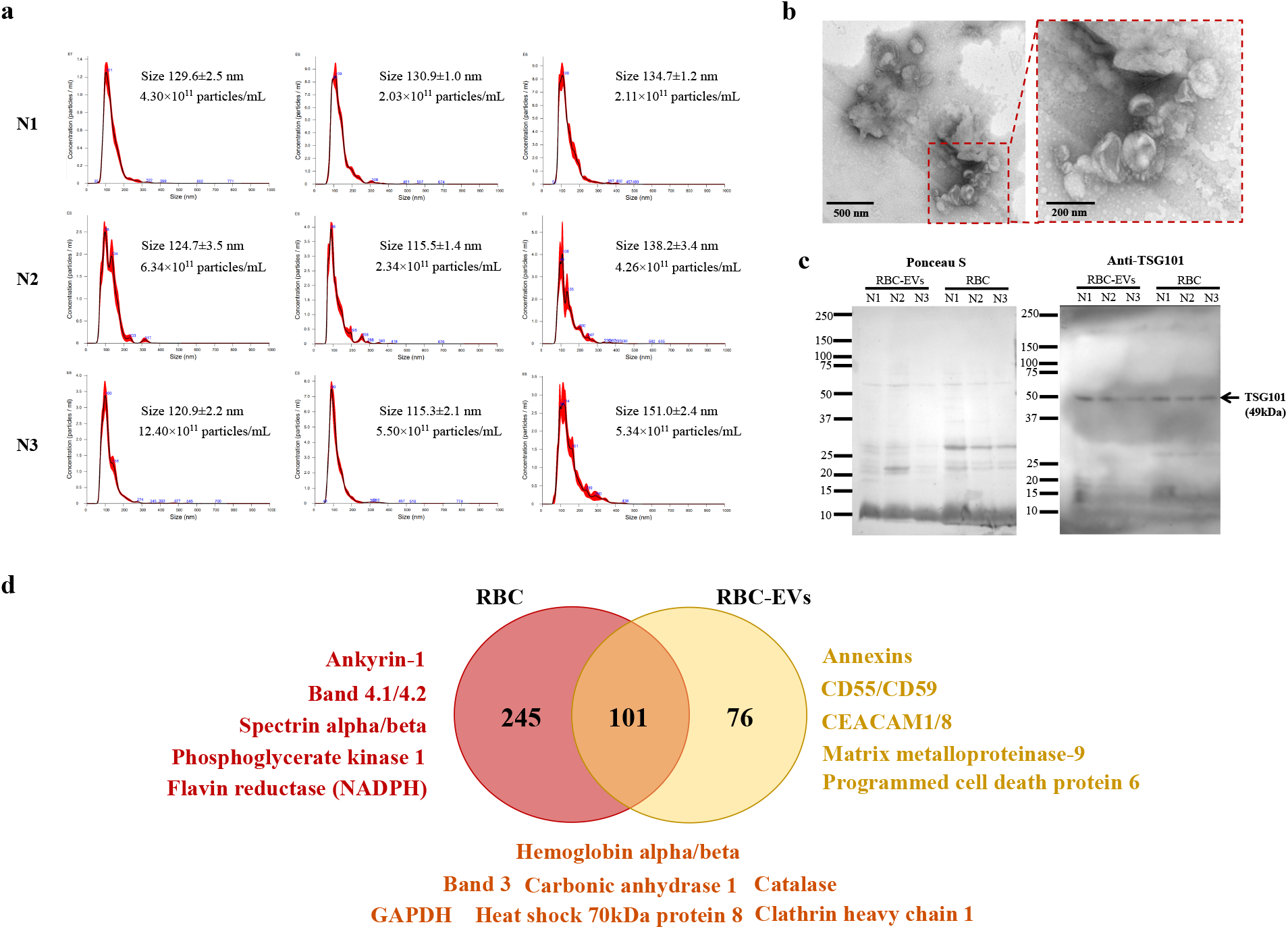
Validation of RBC-EV presence in the isolates. RBC-EVs were produced and isolated from 3 blood group O healthy donors (N1, N2, N3). (**a**) Nanoparticle tracking analysis shows RBC-EV particle size distributions and concentrations. (**b**) Transmission electron microscopy with negative staining exhibits RBC-EVs as round shape nanoscale vesicles. (**c**) TSG101 detection in RBC-EVs by Western blot analysis (20 μg protein/lane); left panel, Ponceau S staining of all protein bands on the electro-transferred membrane; right panel, the full-length blot image shows the EV marker TSG101. (**d**) Venn diagram showing the numbers of common and unique proteins identified from RBC-EVs compared to the originate RBC by mass spectrometric proteomics (full details in **Tables S1-S3**). CEACAM, carcinoembryonic antigen-related cell adhesion molecule; GAPDH, glyceraldehyde-3-phosphate dehydrogenase.

Moreover, the specific EV protein marker in RBC-EVs was confirmed by anti-TSG101 antibody which was expressed at 49 kDa (**Figure 1c**). Mass spectrometric-based proteomics was performed to deliver additional protein evidence of RBC-EVs compared to the originate RBC (**Figure 1d** and **Tables S1-S3**). As expected, hemoglobin was detected in both RBC and RBC-EVs. Glycosylphosphatidylinositol (GPI)-anchored proteins, i.e., CD55 and CD59, plasma membrane-associated protein annexins, and transmembrane protein CEA cell adhesion molecule (CEACAM) 1/8 were uniquely identified in the RBC-EVs. Clathrin heavy chain 1, a protein involved in vesicle trafficking, and cytosolic markers such as heat shock 70kDa protein 8, catalase, and glyceraldehyde-3-phosphate dehydrogenase (GAPDH) were found in both RBC and RBC-EVs (**Figure 1d** and **Tables S1-S3**). These findings indicated the successful production of RBC-EVs, and supported their testing in further experiments.

The basal effect of the native RBC-EVs at doses of 100, 1000 and 10,000 particles/cell was measured on cell viability of multiple cell types, including neuroblastoma cells (*MYCN*-amp: SK-N-BE2; *MYCN*-NA: SH-SY5Y, SK-N-AS, SK-N-SH), and cells derived from lung cancer (A549), breast cancer (MDA-MB-231), liver cancer (HepG2), colon cancer (HT-29), acute T leukemic (Jurkat) cells, and normal cells, e.g., peripheral blood T cells and human fibroblasts (HEK293T) as shown in **Figure 2**. Overall, the native RBC-EVs slightly increased cell viability of SK-N-BE2, SH-SY5Y, MDA-MB-231 and HEK293T cells, but not of SK-N-SH and HepG2, while A549 and HT-29 cells had a trend toward decreased viability. RBC-EVs had no effect on SK-N-AS and T cells, while the native RBC-EVs at 10,000 particles/cell apparently increased cell viability in HEK293T human fibroblasts. These results suggest that the native RBC-EVs may exhibit heterogenous effects on cell viability, depending on the doses and recipient cells, but those effects are small at doses up to 1000 particles/cells. To minimize the basal effect of RBC-EVs on target cells, 1000 RBC-EV particles/cell was used throughout this study.

**Figure 2.**
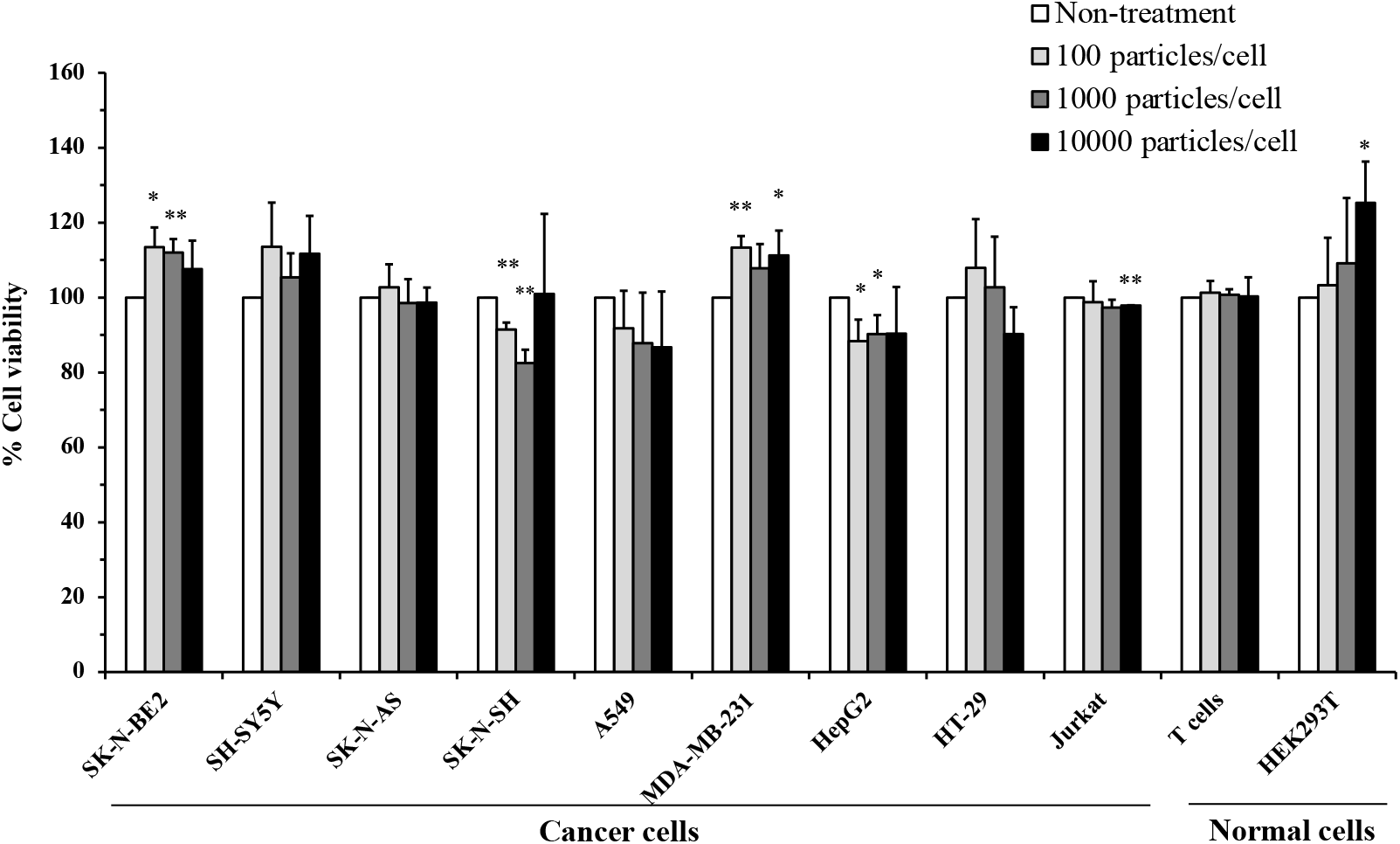
Effect of the native RBC-EVs on cell viability of multiple cell types. Cancer cell lines including neuroblastoma (SK-N-BE2, SH-SY5Y, SK-N-AS and SK-N-SH), lung cancer (A549), breast cancer (MDA-MB-231), liver cancer (HepG2), colon cancer (HT-29), leukemia (Jurkat) and normal cells including T cells and HEK293T cells were treated with native RBC-EVs at varied dosages of 100, 1000 and 10000 particles/cell for 72 hr. Non-treatment served as the control (100% cell viability). \**p*<0.05, \*\**p*<0.01 compared to non-treatment condition.

### 3.3. Optimizing electroporation conditions for miRNA loading into RBC-EVs

Electroporation was tested for its suitability to load miRNA into RBC-EVs. To optimize the electroporation conditions, 5’FAM-tagged miR204 was loaded into RBC-EVs as follows: one pulse for 20ms at 50V, 100V, 250V, 400V and for 1ms at 750V. PKH26 lipophilic dye was used to stain RBC-EV lipid membrane. SK-N-BE2 cells were used as the recipient cells to observe 5’-FAM and PKH26 signal incorporation as representative of the cellular-EV uptake. As shown in **Figure 3a**, the 5’-FAM (green) and PKH26 (red) signals were greater in SK-N-BE2 cells (blue, nuclear staining) when 5’-FAM-tagged miR-204 were loaded into RBC-EVs by the electroporation at the higher voltages of 250V-750V. Unfortunately, the conditions of 400V for 20ms or 750V for 1ms in one pulse caused aggregation and clumping of RBC-EVs; therefore, 250V for 20ms was further tested by the number of pulses (1, 5, 10 times). The higher numbers of pulsing resulted in the higher 5’FAM signals in SK-N-BE2 cells (**Figure 3b**). Accordingly, the electroporation parameters of 250V for 20ms in 10 pulses were used to prepare miRNA-loaded RBC-EVs (100 picomole miRNA in 1x10^9^ RBC-EV particles; details in Methods) for further testing.

**Figure 3.**
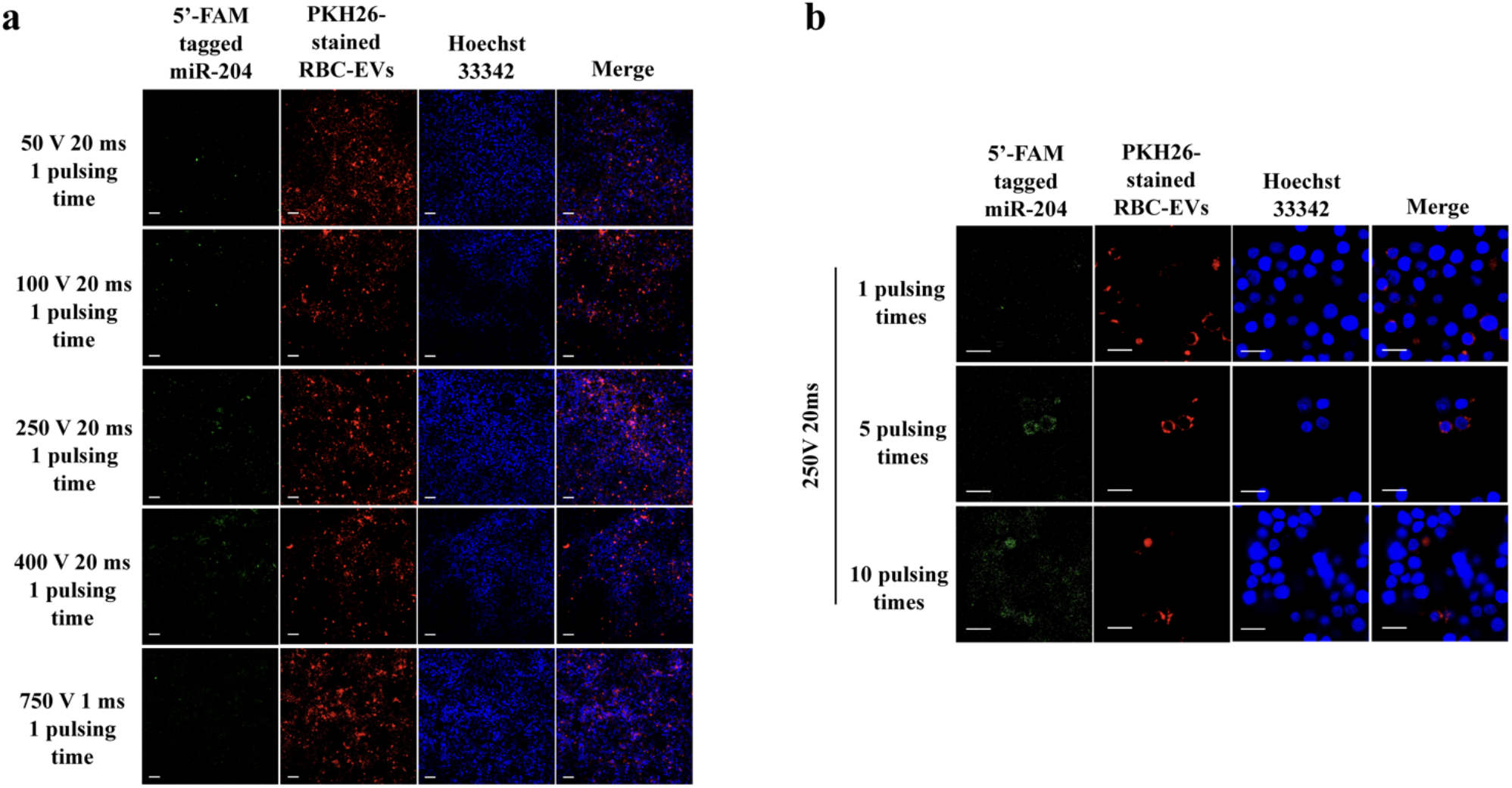
Optimizing electroporation parameters for miRNA loading into RBC-EVs. One hundred picomole of 5’-FAM-tagged miR204 were loaded into 1x10^9^ RBC-EVs with varied electroporation conditions and then incubated with SK-N-BE2 cells (at the dosage of 1000 EV particles/cell) for 2 days. The fluorescence signals from 5’-FAM-tagged miR-204 (green), PKH26-stained RBC-EVs (red) and Hoechst 33342-stained nucleus (blue) were detected by a confocal microscope. (**a**) Varying voltages of 50V, 100V, 250V, 400V for 20 ms or 750V for 1 ms with one pulse. Scale bar, 100 μm. (**b**) Electroporation using 250V for 20ms in 1, 5 and 10 pulses. Scale bar, 20 μm.

### 3.4. RBC-EV^miR204^ repressed MYCN expression in MYCN-amp NB cells

MiR-204 functions as a tumor suppressor in several cancer types including NB [12-16,21,38-41]. *MYCN*, the most important genetic marker in NB, is one of multiple genes targeted by miR-204 [12-20]. We examined whether RBC-EV^miR-204^ could suppress *MYCN* expression in *MYCN*-amp NB cells, with *MYCN*-NA cells used as the comparator (**Figure 4**). Consistent with a previous study [20], our results showed miR-204 in RBC-EVs significantly downregulated *MYCN* expression in SK-N-BE2 cells compared to RBC-EV^miRNC^, the native RBC-EVs and non-treatment conditions, but the magnitude of inhibition was not large (**Figure 4a)**. *MYCN* mRNA levels in *MYCN*-NA SH-SY5Y cells were 100 times less than *MYCN*-amp SK-N-BE2 cells, with no apparent effect of RBC-EV^miR-204^ mediated *MYCN* suppression being observed (**Figure 4b**).

**Figure 4.**
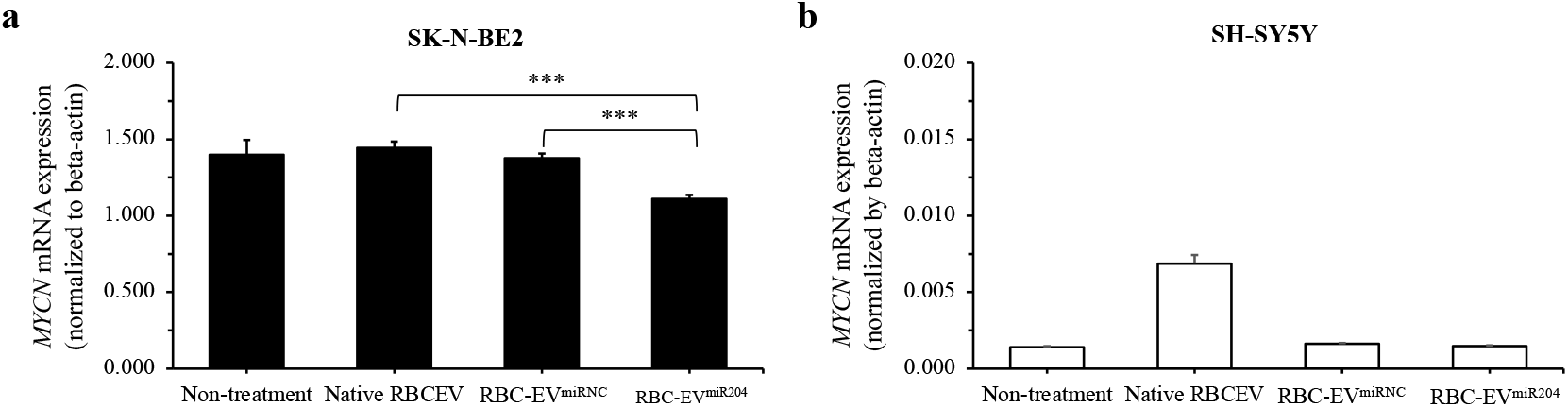
*MYCN* mRNA expression in 2 neuroblastoma cell lines after RBC-EV^miR-204^ treatment. Non-treatment, the native RBC-EVs, RBC-EV^miRNC^ or RBC-EV^miR-204^ at 0.25 µM miRNA loading concentrations treated in SK-N-BE2 and SH-SY5Y cells for 48 h. MYC-N mRNA expression level was measured by qPCR and normalized to actin in both SK-N-BE2 (**a**) and SH-SY5Y (**b**) cells. The statistically significant MYC-N expression was calculated and compared between miRNC and miR204 loaded RBC-EVs treated cells. ***, *p*<0.001.

### 3.5. Effects of RBC-EV^miR204^ on NB cell viability, migration and spheroid formation

Anti-neuroblastoma effects of RBC-EV^miR-204^ were evaluated in *MYCN*-amp SK-N-BE2 and *MYCN*-NA SH-SY5Y NB cells. **Figure 5** demonstrated that RBC-EV^miR-204^ at 1000 particles/cell decreased cell viability of *MYCN*-amp and *MYCN*-NA NB cells relative to those of the native RBC-EVs and non-treatment conditions. Interestingly, RBC-EV^miRNC^ (miRNC loaded RBC-EVs) treatment at 1000 particles/cell also exhibited a cytotoxic effect, but with less potency than RBC-EV^miR-204^ (**Figure 5**).

**Figure 5.**
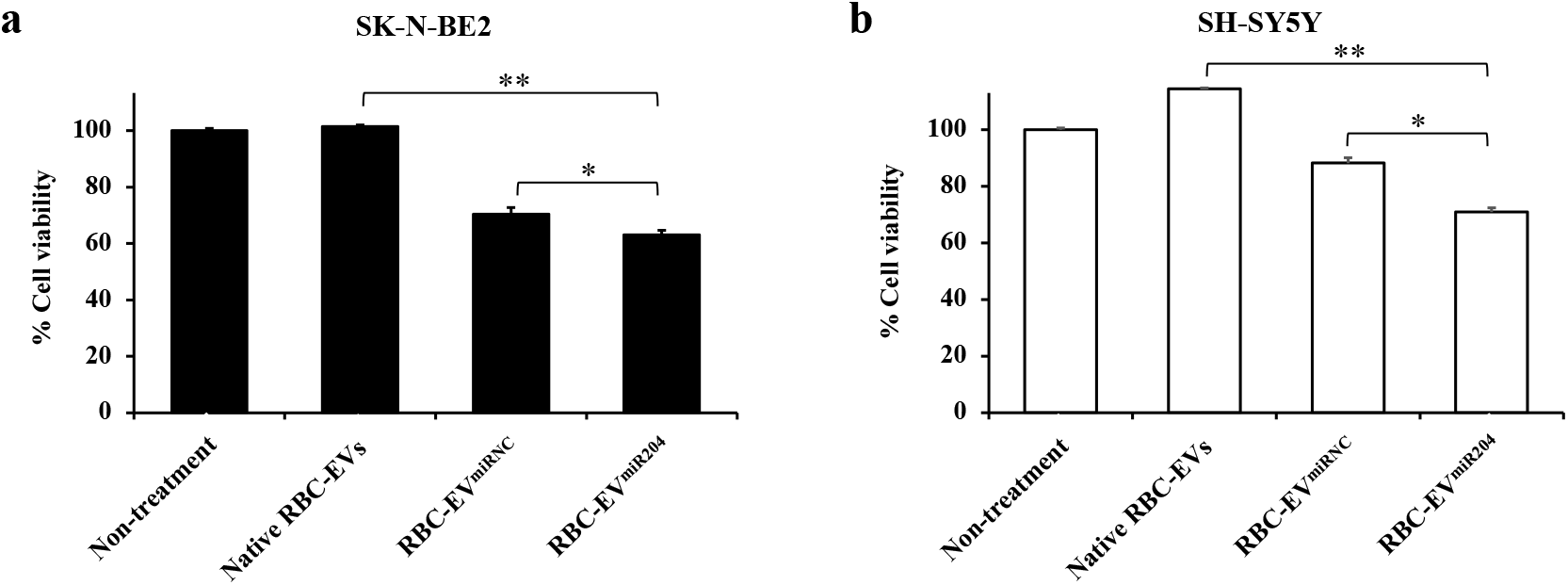
Effects of RBC-EV^miR-204^ on cell viability of *MYCN*-amp and *MYCN*-NA neuroblastoma cells. SK-N-BE2 (*MYCN*-amp NB) and SH-SY5Y (*MYCN*-NA NB) cell lines were treated with the native RBC-EVs, RBC-EV^miRNC^ or RBC-EV^miR-204^ at 1000 particles/cell for 48 hr. Cell viability was detected by the MTT assay. Non-treatment control values were set as 100% cell viability. This experiment was performed in triplicate. *, *p*<0.05, **, *p*<0.01.

Whether RBC-EV^miR-204^ can exhibit anti-NB metastasis was also evaluated by the cell migration-wound scratch assay. Cell migration capability of *MYCN*-amp SK-N-BE2 cells were markedly decreased after RBC-EV^miR-204^ treatment compared to the native RBC-EVs and non-treatment conditions, while RBC-EV^miRNC^ exhibited a moderate effect against SK-N-BE2 cell migration (**Figure 6a**). In SH-SY5Y cells, however, no obvious anti-migration activity was observed with either RBC-EV^miR-204^, RBC-EV^miRNC^ or the control conditions (**Figure 6b**). Note that the innate cell migration capability of SH-SY5Y cells (the representative of *MYCN*-NA NB) is much lower than SK-N-BE2 cells (the representative of *MYCN*-amp high-risk NB).

**Figure 6.**
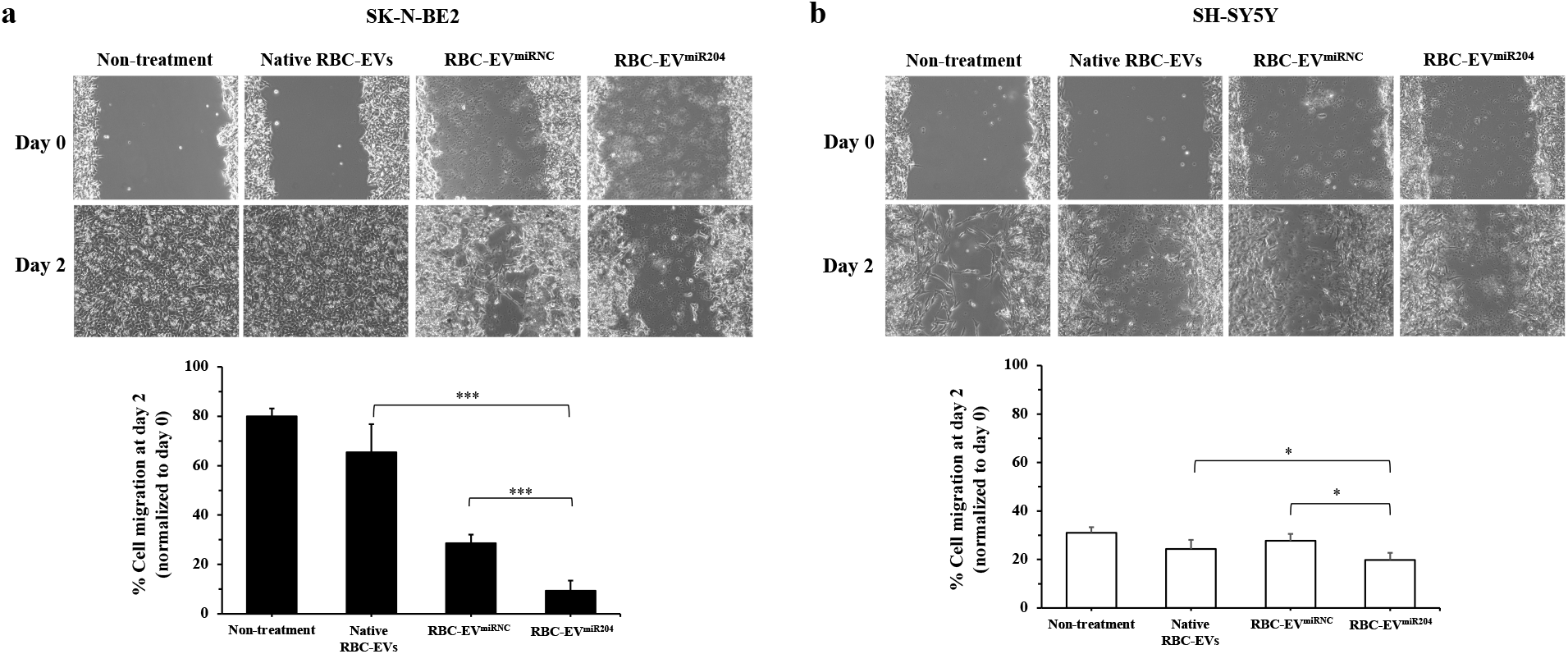
Effects of RBC-EV^miR-204^ on cell migration of *MYCN*-amp and *MYCN*-NA neuroblastoma cells. After reaching 100% confluence, a 200-μL pipette tip was used to create wound onto *MYCN*-amp SK-N-BE2 (**a**) and *MYCN*-NA SH-SY5Y (**b**) cells. Thereafter, cells were treated with the native RBC-EVs, RBC-EV^miRNC^ or RBC-EV^miR-204^ at 1000 particles/cell. The non-treatment condition served as the blank control. The wound width at day 2 was measured and converted to the percentage of cell migration as normalized by the wound width at day 0. *, p<0.05; ***p<0.001.

Following migration, tumor spheroid formation is a crucial process in the progression of metastasis [42]. By applying ultra-low attachment microplates, neuroblastoma cell lines can form 3D spheroids *in vitro* within 24 h (see Methods). Four neuroblastoma spheroids of *MYCN*-amp (SK-N-BE2 and SK-N-DZ) (**Figure 7a,b**) and *MYCN*-NA (SH-SY5Y and SK-N-SH) NB cells (**Figure 7c,d**) were formed in the presence or absence of RBC-EV^miR-204^ at 1000 particles/cell. RBC-EV^miR-204^ significantly suppressed NB spheroid formation and growth, regardless of *MYCN* amplification, relative to RBC-EV^miRNC^, native RBC-EVs, or non-treatment conditions. Overall, RBC-EV^miR-204^ at 1000 particles/cell can exert anti-neuroblastoma effects *in vitro*.

**Figure 7.**
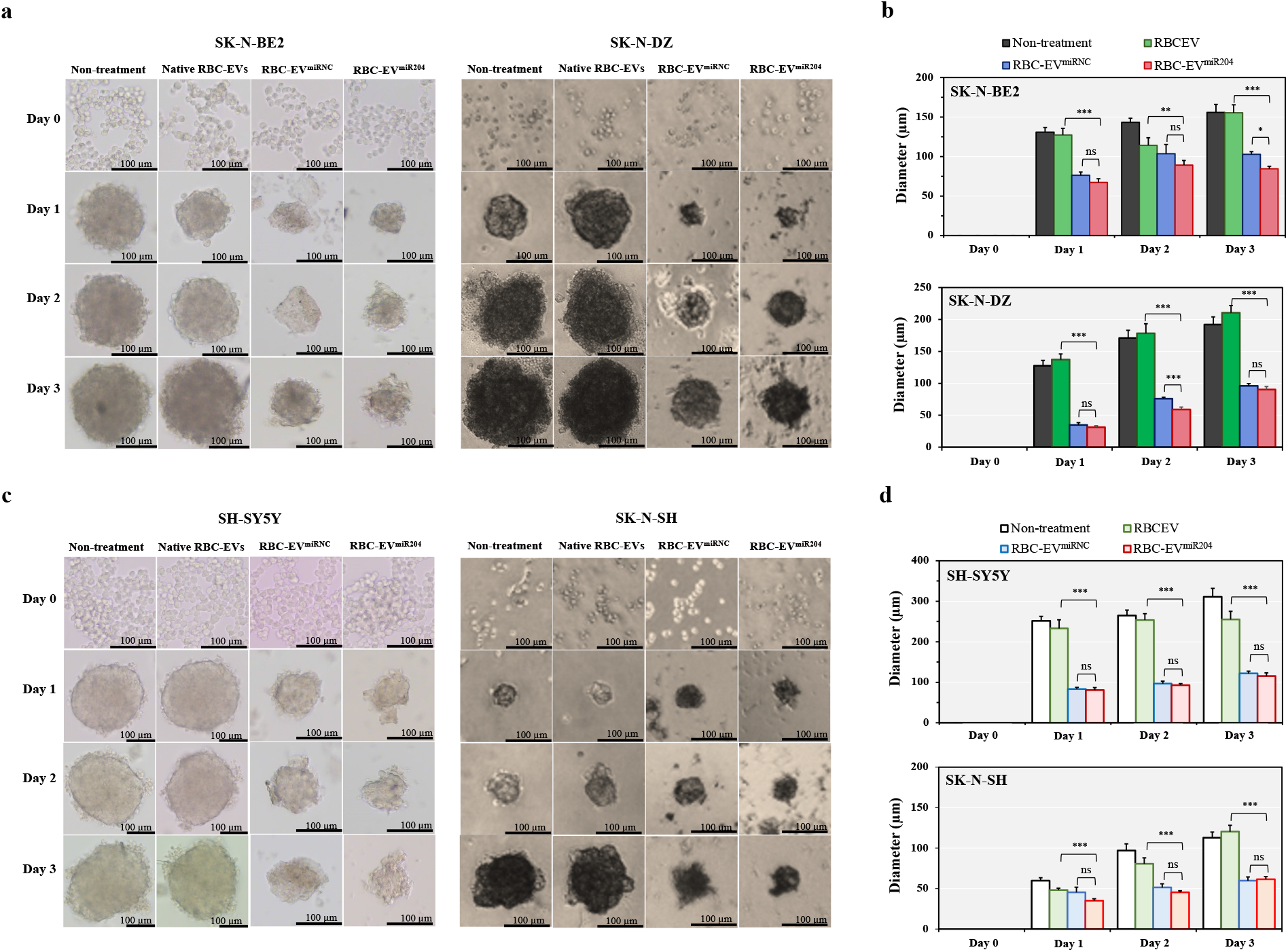
Effects of RBC-EV^miR-204^ on spheroid formation and growth in four distinct neuroblastoma cell lines: SK-N-BE2, SK-N-DZ (*MYCN*-amp), SH-SY5Y, and SK-N-SH (*MYCN*-NA). NB cells were mixed with the native RBC-EVs, RBC-EV^miRNC^ or RBC-EV^miR-204^ at 1000 particles/cell and were allowed to form spheroids for 3 days (details in the Methods section). (**a**) Representative images of spheroid formation and growth from day 0 to day 3, and (**b**) spheroid diameter measured by ImageJ software of *MYCN*-amp NB cells. (**c**) The representative images of spheroid formation and growth from day 0 to day 3, and (**d**) spheroid diameter measured by ImageJ software of MYCN-NA cells. The non-treated cells served as the blank control. This experiment was performed in three biological replicates. ns, not significant; *, *p*<0.05; **, *p*<0.01; ***, *p*<0.001.

### 3.6. SWATH-proteomics detected modes of action of RBC-EV^miR-204^ against neuroblastoma

In fact, miRNAs target multiple genes to exert biological effects [43]. While miR-204 can repress *MYCN* expression (**Figure 4**), RBC-EV^miR-204^ can exhibit anti-neuroblastoma effects regardless of *MYCN* status (**Figures 5-7**). These findings support a hypothesis that RBC-EV^miR-204^ trigger (unidentified) genes/pathways that play crucial roles in NB tumorigenesis and progression.

To elucidate the complex molecular/pathway alterations in response of anti-neuroblastoma effects of RBC-EV^miR-204^ in both *MYCN*-amp and *MYCN*-NA NB cells, SK-N-BE2 and SH-SY5Y cells treated with RBC-EV^miR-204^, RBC-EV^miRNC^, the native RBC-EVs or non-treatment conditions were subjected to targeted label-free SWATH-proteomics operating on a Triple-TOF 6600+ mass spectrometer (Methods). Accordingly, 1216 and 1111 unique peptides were detected and quantified at FDR<0.01 in SK-N-BE2 (**Table S4**) and SH-SY5Y treatment groups (**Table S5**), respectively. After multiple comparison, 170 and 176 significantly altered proteins were detected in RBC-EV^miR-204^ treated SK-N-BE2 (**Table S6**) and SH-SY5Y (**Table S7**), respectively compared to RBC-EV^miRNC^, the native RBC-EVs and non-treatment conditions. A Venn diagram illustrates the common altered proteins and the unique changes in SK-N-BE2 and SH-SY5Y cells. As shown in **Figure 8**, 60 proteins were altered in both cell lines, while 110 and 116 proteins were uniquely altered in SK-N-BE2 and SH-SY5Y cells, respectively (**Table S8**). STRING protein-protein interaction and Reactome pathway enrichment analysis revealed that alterations in RNA metabolism was the top enriched pathway in RBC-EV^miR204^ treated *MYCN*-amp SK-N-BE2 cells, but not in *MYCN*-NA cells (**Figure 8** and **Table S9)**. The proteins altered in common across both types of cells were also significantly matched to RNA metabolism following RBC-EV^miR-204^ treatment (**Figure 8** and **Table S9)**. Dysregulation of ribosome biogenesis has been recognized as a new therapeutic target of *MYCN*-amp neuroblastoma [44,45]. Our findings reinforce the conclusion that RBC-EV^miR-204^ mediated ribosome dysregulation as a major mechanism of action against pediatric neuroblastoma.

**Figure 8.**
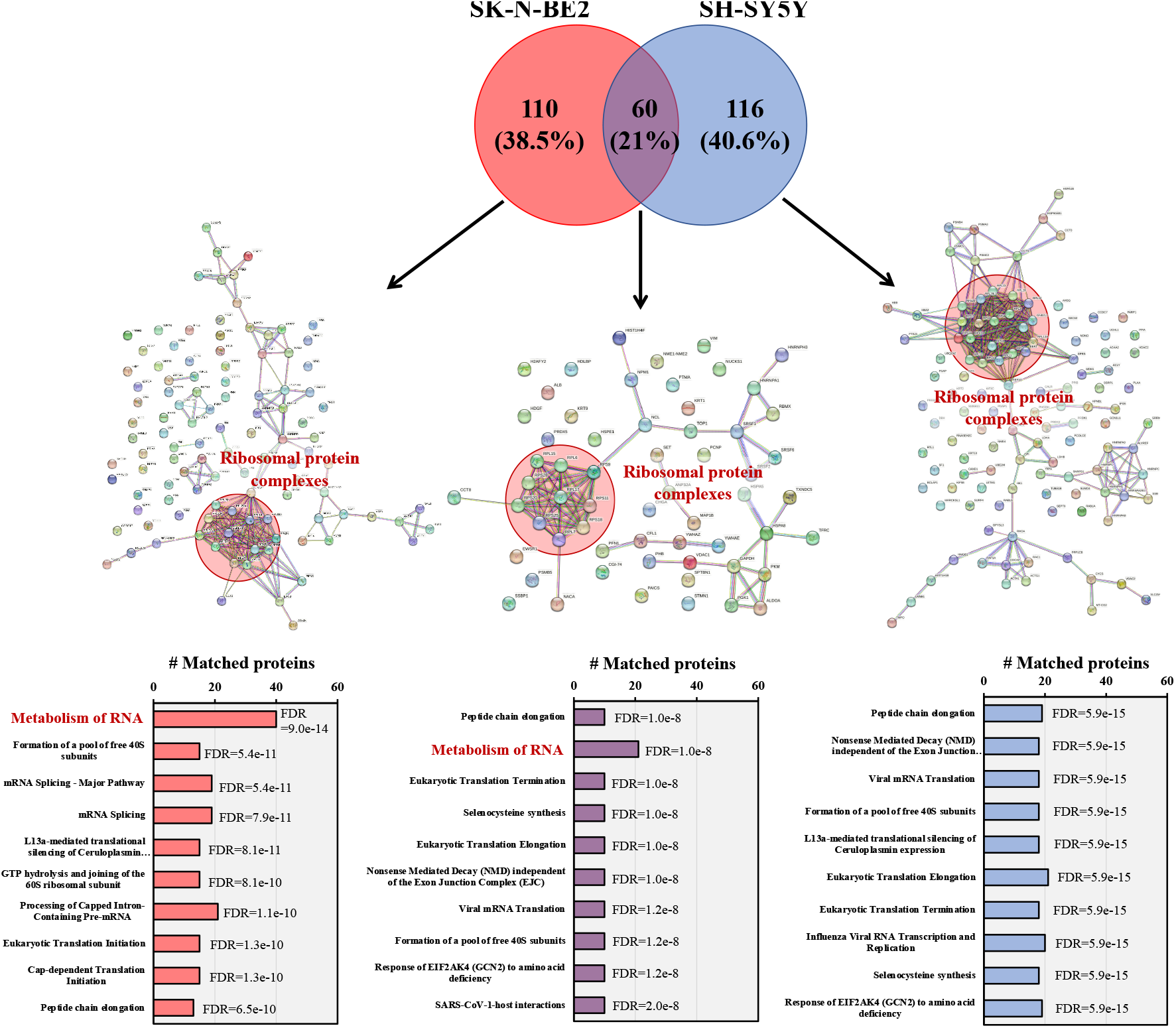
SWATH-proteomics and bioinformatics to unravel the mode of action of RBC-EV^miR-204^ in neuroblastoma cells. Differentially expressed proteins were obtained from SWATH-proteomics of SK-N-BE2 and SH-SY5Y cells treated with RBC-EV^miR-204^ compared to RBC-EV^miRNC^, native RBC-EVs and non-treatment conditions (full data in **Tables S4-S7**) and expressed as a Venn diagram, STRING protein interaction and Reactome enrichment (details in **Tables S8**,**S9**). Alterations in RNA metabolism and ribosome dysregulation are, at least in part, responsible for the anti-neuroblastoma effects of RBC-EV^miR-204^. FDR, false discovery rate.

## 4. Discussion

High-risk pediatric neuroblastoma requires intensive multimodal therapy including induction chemotherapy, surgical resection, autologous stem cell transplantation, radiation, immunotherapy, and cytokines plus isotretinoin [8,44]. Despite this intensive multimodal regimen, suitable therapeutic efficacies have not been achieved. New treatment modalities are needed. Herein, a novel EV-in treatment strategy [46] is proposed to utilize RBC-EV^miR-204^ as a new mode of adjunct therapy against high-risk neuroblastoma.

To load miRNAs into RBC-EVs by electroporation, the electroporation settings were examined for their effects upon cells (**Figures S1 and S2**) and then RBC-EVs (**Figure 3**). The low voltage can load lower quantities of miRNAs into RBC-EVs compared to the high voltage, but preserves the EV cargo delivery capacity (**Figure 3a**). Higher voltage, even with a brief period of time (1 ms), caused protein and miRNA denaturation and breakdown. A combination of lower voltage (250 V) for 20 ms as 10 pulses improved miRNA loading into RBC-EVs while preserving the essential EV properties (**Figure 3b**), thus defining the optimized electroporation protocol for RBC-EV^miR-204^ production.

After optimizing electroporation parameters for loading miRNA to RBC-EVs, anti-neuroblastoma effects of RBC-EV^miR-204^ were evaluated in both *MYCN*-amp and *MYCN*-NA neuroblastoma cells by using cell viability, cell migration and spheroid formation and growth assays. RBC-EV^miR-204^ decreased cell viability, mitigated cell migration, and suppressed spheroid formation and growth, more strongly against *MYCN*-amp NB cells, but these anti-neuroblastoma effects were also observed, but to a lesser extent, in *MYCN*-NA cells (**Figures 5-7**). These results are consistent with previous evidence, in which miR-204 exhibited anticancer activities against several cancers with diversed oncogenic backgrounds, i.e., oral squamous cell carcinoma (OSCC), renal cell carcinoma, and non-small-cell lung cancer [47-50]. As RBC-EV^miR-204^ could inhibit both NB cell migration and spheroid formation and growth, the use of RBC-EV^miR-204^ as an adjuvant therapy may mitigate metastasis in high-risk neuroblastoma, regardless *MYCN* amplification status.

The anti-neuroblastoma effects of RBC-EV^miR204^ do not seem to be fully explained by *MYCN* suppression (**Figure 4**), and effects were also observed on *MYCN*-NA cells (**Figures 5-7**). SWATH-proteomics was conducted to study the complex underlying mechanisms of RBC-EV^miR-204^ against both *MYCN*-amp and *MYCN*-NA NB cells (**Figure 8**). Alterations in RNA metabolism and ribosome dysfunction may be responsible for the major therapeutic action of RBC-EV^miR-204^ in both *MYCN*-amp and *MYCN*-NA NB cells. Dysregulation of ribosome biogenesis has been implicated as a new target for treating *MYCN*-amp neuroblastoma [44,45]. However, living cells need bioenergy and biosynthesis in cellular metabolism, and cancers, regardless of type, need higher protein levels in metabolic pathways to promote their rapid cell growth and proliferation [51]. In this regard, RBC-EV^miR-204^ may have broad anticancer effects against many cancer types, and therefore warrant further investigation in preclinical models and clinical studies.

There are several limitations to this study: First, while this study used confocal microscopy to detect co-localization of 5’-FAM-tagged miR-204 and PKH26-stainsed RBC-EVs onto neuroblastoma cells as evidence of cellular-EV uptake, the loading efficiency of miR-204 into RBC-EVs, as well as the releasing efficiency of miR-204 cargo into the recipient cells, could not be determined. Single-molecule localization-based super-resolution microscopy should be used to address this issue [52,53].

Second, while this study successfully demonstrated anti-neuroblastoma activities of RBC-EV^miR-204^ *in vitro*, factors affecting pharmacokinetics of RBC-EV^miR-204^ should be determined before investigating therapeutic effects *in vivo*. For example, the systemic delivery of EVs, including RBC-EVs, resulted in a rapid systemic clearance of EVs by phagocytes at liver and spleen. Other routes of EV administration, i.e., subcutaneous injection or oral gavage, reduced liver and spleen accumulation and exhibited a broader biodistribution toward multiple tissues, including gastrointestinal tract, pancreas, kidney, heart, lungs, and brain [54-56].

Third, while this study focused on *MYCN* as the primary target of miR-204, multiple gene targets of miR-204 [12-14,16,18,38-40] may allow RBC-EV^miR-204^ to exhibit anticancer effects against broader types of malignancies; this could be tested.

Fourth, RBC-EV^miRNC^ tend to inhibit neuroblastoma cells, although at less potency than RBC-EV^miR-204^. MiRNC mimic sequence is derived from *Caenorhabditis elegans* with no homology to human and mouse gene sequences, has not been identified as a negative regulator of neuroblastoma, or as having any effect on *MYCN* gene (**Figure 4**). While exploring this unexpected finding of RBC-EV^miRNC^ was beyond the focus of this study, gene target prediction (**Table S10**) by miRDB (www.mirdb.org) revealed HDAC1 as this miRNC (5’-CGGUACGAUCGCGGCGGGAUAUC-3’) may interact with HDAC1, a potential therapeutic target of NB [57,58]. Future investigations on the specific targets and mechanisms of action of the miRNC, either alone or with carrier loading, could further extend the translational potential of miRNA-based therapeutics, particularly for high-risk neuroblastoma.

## 5. Conclusions

This study utilized RBC-EVs as a biocompatible carrier of miR-204 mimics and demonstrated anticancer effects of RBC-EV^miR-204^ in *MYCN*-amp and *MYCN*-NA neuroblastoma cells. RBC-EV^miR204^ induced alterations in RNA metabolism and induced ribosome dysregulation, thereby suppressing cell viability, migration, spheroid formation, and growth. Future investigations should evaluate therapeutic effects of RBC-EV^miR-204^ in preclinical models and clinical studies.

## Acknowledgments

We would like to thank Dr. Supasek Kongsomros and Dr. Tassanee Lerksuthirat to technical support. Furthermore, we thank to staffs at Research Center and Central Laboratory of Pediatric Department at Faculty of Medicine Ramathibodi Hospital to facilitate materials, equipment and space.

## Author Contributions

Conceptualization, W.C. and S.C.; Methodology, W.C., J.P. and S.K.; Software, W.C. and S.C.; Validation, W.C., D.S.N., A.L.M., S.H. and S.C.; Formal Analysis, W.C., J.P., S.K.; Investigation, W.C., J.P., S.K.; Resources, W.C., D.S.N., A.L.M., S.H.; Data Curation, W.C. and S.C.; Writing – Original Draft Preparation, W.C.; Writing – Review & Editing, W.C., J.P., S.K., D.S.N., A.L.M, S.H. and S.C.; Visualization, W.C. and S.C.; Supervision, S.H. and S.C.; Project Administration, W.C.; Funding Acquisition, W.C. and S.H. All authors have read and agreed to the published version of the manuscript.

## Data Availability Statement

All data are available in the main text or the supplementary materials.

## Supplementary tables

Table S1. A list of 346 identified proteins in RBC by mass spectrometric proteomics.

Table S2. A list of 177 identified proteins in RBC-EVs by mass spectrometric proteomics.

Table S3. Lists of UniProt ID of commonly and uniquely identified proteins in RBC and RBC-EVs by mass spectrometric proteomics.

Table S4. Raw expression data of SWATH-proteomics of 4 treatment groups in MYCN-amp SK-N-BE2 cells.

Table S5. Raw expression data of SWATH-proteomics of 4 treatment groups in MYCN-non-amp SH-SY5Y cells.

Table S6. Altered expressed proteins among 4 treatment groups in MYCN-amp SK-N-BE2 cells.

Table S7. Altered expressed proteins among 4 treatment groups in MYCN-non-amp SH-SY5Y cells.

Table S8. Lists of UniProt ID of altered expressed proteins from SWATH-proteomics for Venn diagram, STRING protein network and Reactome pathway enrichment analyses.

Table S9. Top 10 significantly enriched Reactome pathways of the differentially expressed proteins from 4 treatment groups in SK-N-BE2 and SH-SY5Y cells.

Table S10. Predicted gene targets of miRNC (5’-CGGUACGAUCGCGGCGGGAUAUC-3’) by miRDB (www.mirdb.org).

## Supplementary figures

Figure S1. Vary electroporation parameter for internalizing miRNA to SK-N-BE2 cells.

Figure S2. Vary electroporation parameter for internalizing miRNA to SH-SY5Y cells.

Figure S3. %Cell viability after miR204 internalized into neuroblastoma cells.

Figure S4. %Cell viability of free miR204 treated neuroblastoma cells.

Figure S5. %Cell viability of miR204 transporting using lipofectamine delivery system to SK-N-BE2 cells.

