## Supplementary tables and figures for "Red blood cell-derived extracellular vesicles with miR-204 mimic loading for pediatric neuroblastoma treatment": Supplementary figures_preprint.docx

**SUPPLEMENTARY MATERIALS**

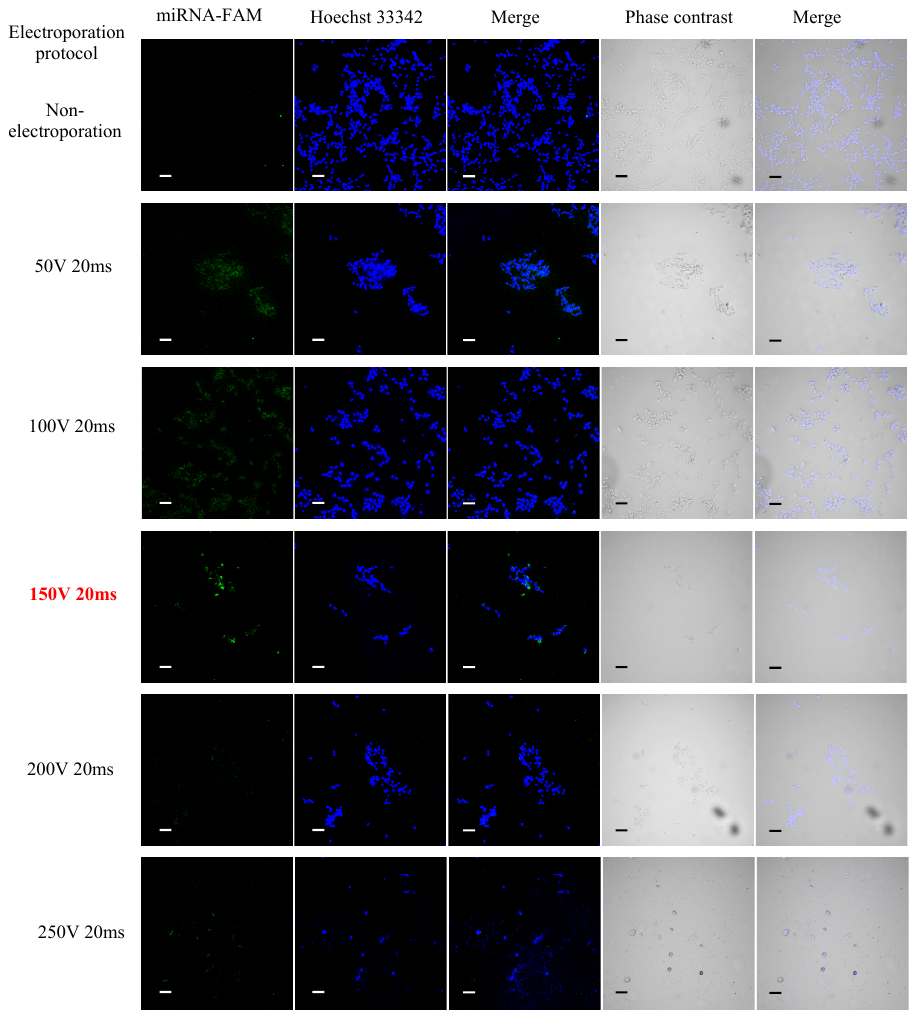

**Figure S1. Evaluation of electroporation parameters for internalizing miRNA to SK-N-BE2 cells.** 2×10^5^ SK-N-BE2 cells were mixed with 100 pmol 5’FAM tagged miRNA and then electroporated using vary parameter i.e., 50V 20ms, 100V 20ms, 150V 20ms, 200V 20ms and 250V 20ms with once pulsing time and without electroporation. The 5’FAM tagged miRNA internalized SK-N-BE2 cells were cultured for 24 h before imaging using confocal microscope. The highest green fluorescent signal was at 150V 20ms. Hoechst 33342 nuclear staining is blue. 200× magnification. Scale bar, 100 μm.

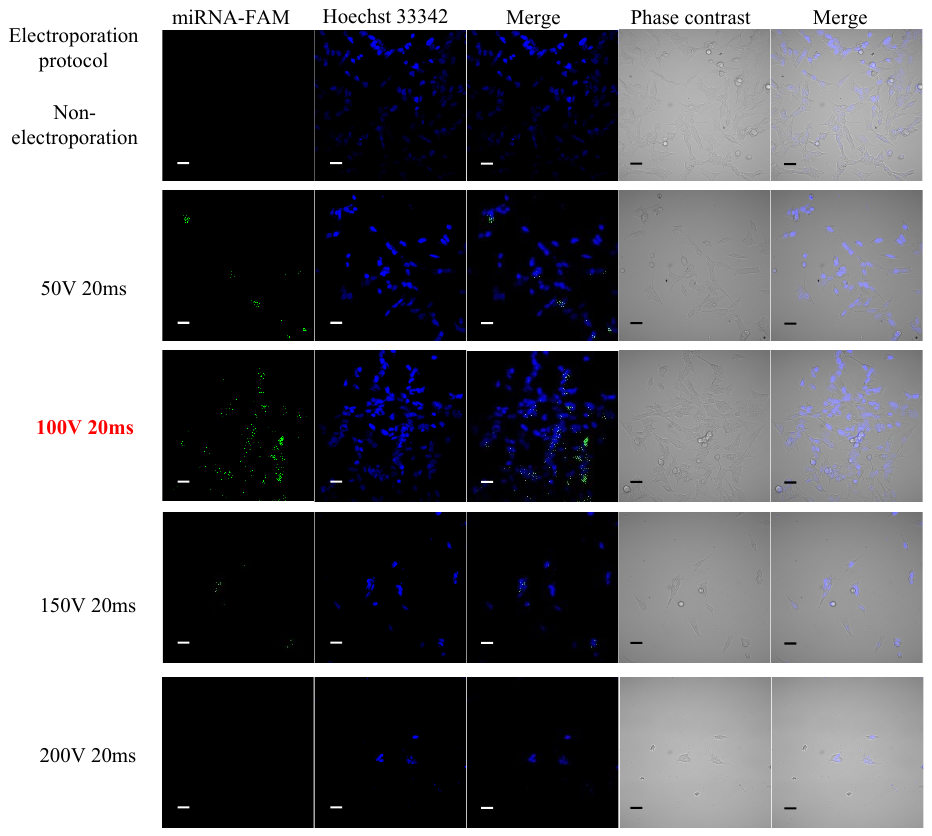

**Figure S2. Evaluation of electroporation parameters for internalizing miRNA to SH-SY5Y cells.** 2×10^5^ SH-SY5Y cells were mixed with 100 pmol 5’FAM tagged miRNA and then electroporated using vary parameter i.e., 50V 20ms, 100V 20ms, 150V 20ms and 200V 20ms with once pulsing time and without electroporation. The FAM tagged miRNA internalized SK-N-BE2 cells were cultured for 24 h before imaging using confocal microscope. The highest green fluorescent signal was at 100V 20ms. Hoechst 33342 nuclear staining is blue. 400× magnification. Scale bar, 50 μm.

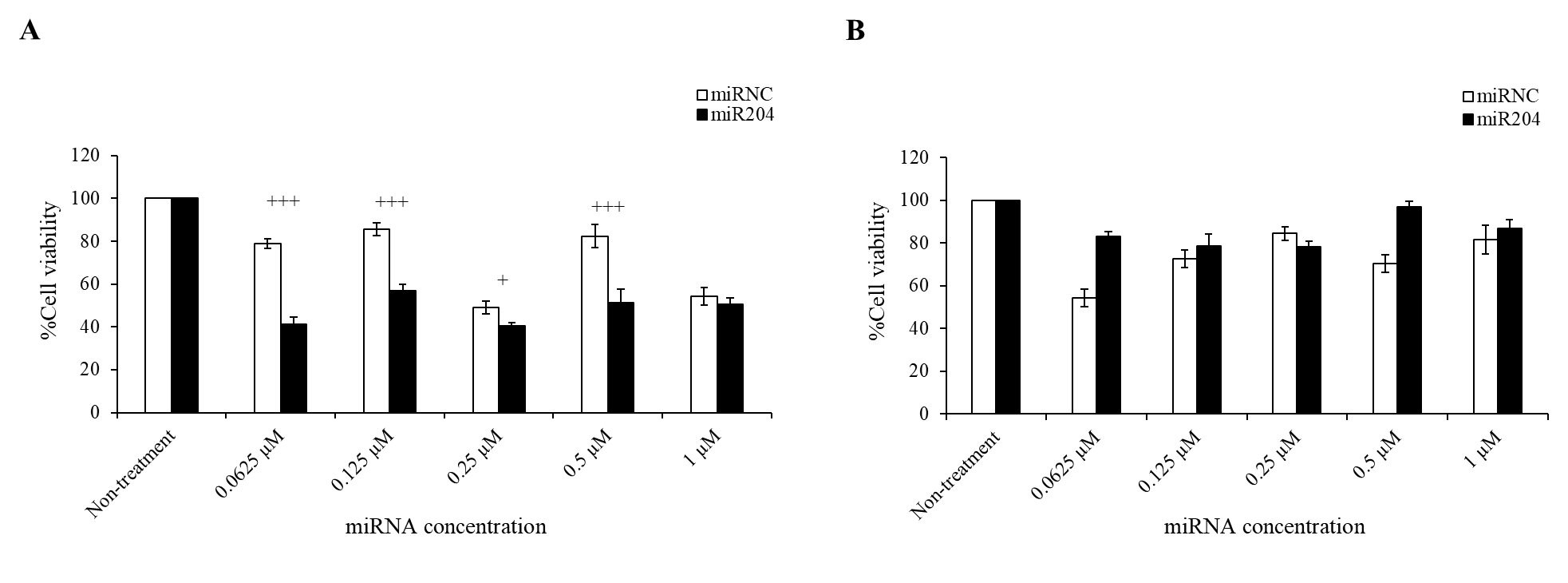

**Figure S3. %Cell viability after miR204 internalized into neuroblastoma cells using electroporation.** SK-N-BE2 (**A**) and SH-SY5Y (**B**) cells were electroporated at 150V 20ms and 100V 20ms, respectively to internalize miRNA at vary concentration i.e., 0-1 µM and then further cultured for 24 h. The %cell viability was measured by MTT assay. + p<0.05, +++p<0.01 were significant difference between miR204 and miRNC treatment (n=3 biological replication).

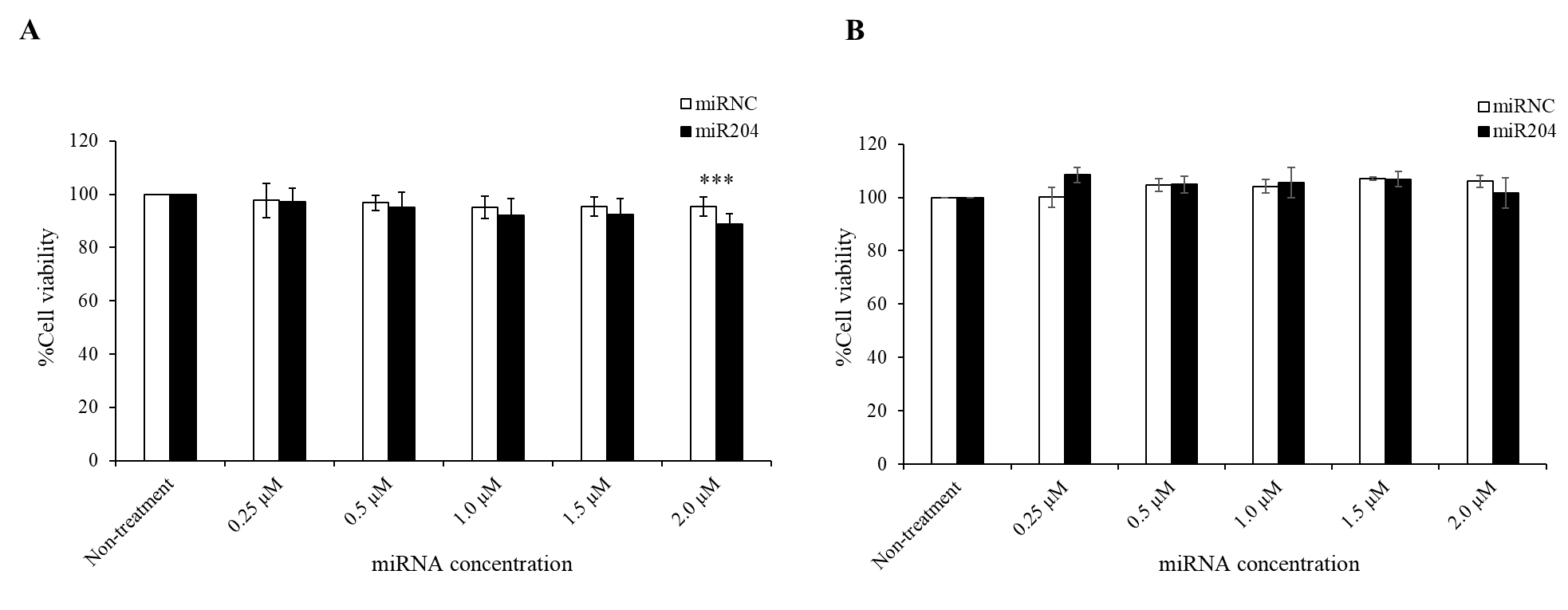

**Figure S4. %Cell viability of free miR204 treated neuroblastoma cells.** Two neuroblastoma cell lines i.e., SK-N-BE2 (**A**) and SH-SY5Y (**B**) cells were treated with free miRNC and free miR204 at vary concentration (0-2 µM) in the medium supplement with 1%FBS for 24 h. The cell viability was measured by MTT assay (mean±SD). ***p<0.01 compared to non-treatment group.

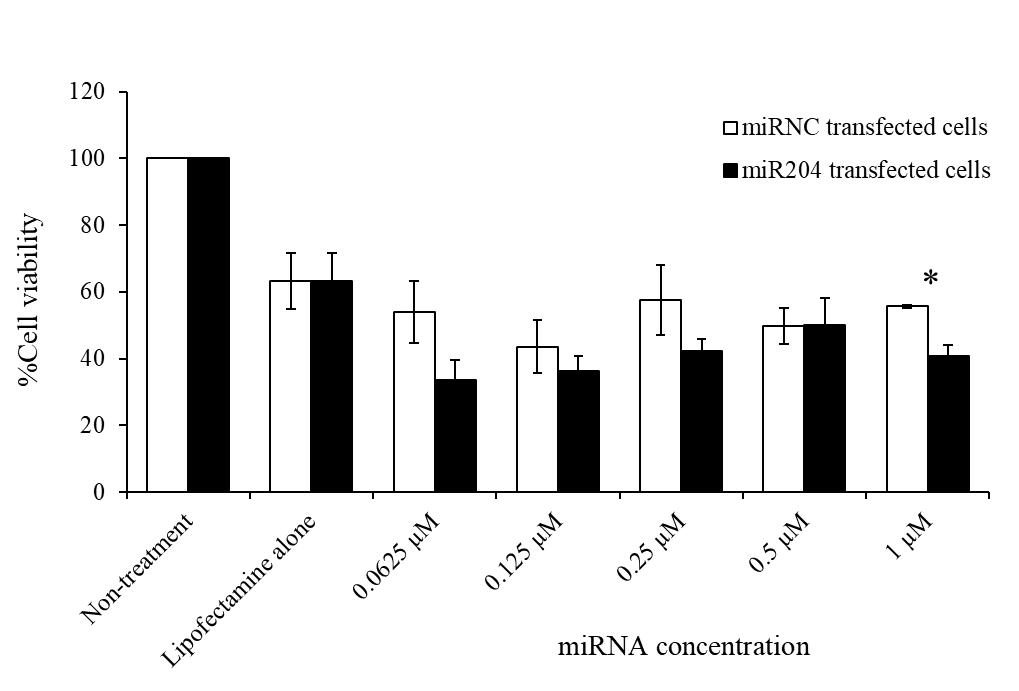

**Figure S5. %Cell viability of miR204 transfection using lipofectamine delivery system in SK-N-BE2 cells.** Lipofectamine was delivery system to transport miRNA at vary concentration (0-1 µM) to neuroblastoma cells and then incubated for 24 h before measuring cell viability using MTT assay (mean±SEM). *p<0.05 comparing to that miRNC treatment.
